# Acetylcholine release from basal forebrain promotes cortical synaptic plasticity via somatostatin interneurons

**DOI:** 10.1101/2023.07.05.547662

**Authors:** Xueyang Zhao, Yu E. Zhang, Lu Yang, Yang Yang

**Affiliations:** School of Life Science and Technology, ShanghaiTech University, Shanghai 201210, China; Department of Neurobiology, Department of Neurosciences, and Center for Neural Circuits and Behavior, University of California San Diego, La Jolla, CA 92093, USA; Department of Biology, University of Virginia, Charlottesville, VA 22904, USA

## Abstract

Basal forebrain (BF) cholinergic projections densely innervate the neocortex, releasing acetylcholine (ACh) to modulate neuronal excitability, thereby influencing sensory processing and cognitive functions. However, the synaptic basis of such ACh-induced effects *in vivo* remains elusive. To investigate how ACh release from BF regulates synaptic dynamics in the neocortex, we selectively stimulated BF cholinergic neurons in mice, and used two-photon microscopy to track structural changes of dendritic spines on cortical pyramidal neurons. We found that optogenetic and chemogenetic stimulation of BF cholinergic neurons rapidly induced spine formation in layer 5 pyramidal neurons in the auditory cortex and the posterior parietal cortex. Selective blockage of ACh receptors (AChRs) demonstrated that nicotinic AChRs are required for the ACh-induced spine formation. We also found that Ach-induced spine formation of pyramidal neurons requires the activity of cortical somatostatin-expressing inhibitory interneurons. Together, our study revealed ACh-induced synaptic plasticity in the cortical microcircuit *in vivo*, providing a synaptic mechanism for cortical neuronal plasticity and functional changes associated with BF ACh release.

## Introduction

The cholinergic system has been associated with a wide range of brain functions (Parikh, Kozak, Martinez, & Sarter, 2007; Saper, Fuller, Pedersen, Lu, & Scammell, 2010; Teles-Grilo Ruivo et al., 2017). Previous studies have reported that during arousal, learning, and attention, basal forebrain (BF) cholinergic neurons release acetylcholine (ACh) in the neocortex, which modulates the activity of cortical neurons (Kim et al., 2016), thereby modifying sensory processing in the sensory cortex (Froemke et al., 2013; Meir, Katz, & Lampl, 2018), and attentional processing in the posterior parietal cortex (PPC) (Bucci, Holland, & Gallagher, 1998). Application of acetylcholine or its agonists, as well as electrical and optogenetic stimulation of the BF, have also been shown to promote neuronal excitability in sensory and higher-order cortices, and influence cortical processing (Froemke et al., 2013; Kilgard & Merzenich, 1998; Weinberger, 2007).

It is well-established that neuronal activity and excitability relies fundamentally on synaptic plasticity (Froemke, 2015). ACh has been implicated in modulating the synaptic plasticity of multiple types of neurons, including glutamatergic pyramidal neurons (PNs) (De Bartolo, Gelfo, Mandolesi, Foti, Cutuli, & Petrosini, 2009) and GABAergic interneurons (Pancotti & Topolnik, 2022), through two main types of cholinergic receptors: nicotinic AChRs (nAChRs) and muscarinic AChRs (mAChRs) (Colangelo, Shichkova, Keller, Markram, & Ramaswamy, 2019). Spines are postsynaptic structures of excitatory neurons, and changes in spine morphology and density are a key component underlying functional changes in neurons (Holtmaat & Svoboda, 2009; Kasai, Fukuda, Watanabe, Hayashi-Takagi, & Noguchi, 2010; Yuste, 2011). Activation of nAChRs can induce the enlargement of spine heads and increase spine number and density in cultured hippocampal neurons (Lozada et al., 2012a, 2012b; Oda et al., 2014). Activation of mAChRs leads to reversible morphological changes in spine heads (Schatzle et al., 2011). These studies suggest a link between ACh and synaptic plasticity, but direct evidence for such effects *in vivo* is still lacking.

Two-photon microscopy has been widely used to study the structural plasticity of synapses *in vivo*, as it can support high-resolution imaging over extended periods with low photo-toxicity, enabling observation of synaptic structural changes, including those of spines and axon terminals, in live animals (Holtmaat et al., 2009; Knott, Holtmaat, Wilbrecht, Welker, & Svoboda, 2006; Trachtenberg et al., 2002). A previous study of the dopaminergic system reported that inhibition of dopamine receptor D1 facilitates spine elimination, whereas inhibition of dopamine receptor D2 promotes spine formation in the mouse motor cortex (Guo et al., 2015), suggesting that neuromodulators can influence cortical synaptic dynamics *in vivo*.

Aiming to characterize the synaptic basis of ACh-induced plasticity in the neocortex *in vivo*, we examined how ACh mediates the structural plasticity of synapses by selectively stimulating cholinergic neurons in the BF and used two-photon microscopy to monitor structural changes in dendritic spines of pyramidal neurons in the auditory cortex (ACx) and posterior parietal cortex (PPC). Briefly, we found that ACh release from BF cholinergic neurons significantly increased spine formation in both cortical areas, and that ACh-induced spine formation requires nAChRs, GABA transmission, and the activity of somatostatin-expressing (SST) inhibitory interneurons.

## Results

### ACh release by optogenetic stimulation of basal forebrain cholinergic neurons induces spine formation in the auditory cortex

BF cholinergic neurons send dense projections to the auditory cortex (ACx) (Mesulam, Mufson, Wainer, & Levey, 1983), and ACh release is essential for regulating sensory processing and plasticity in ACx (Bakin & Weinberger, 1996; Bjordahl, Dimyan, & Weinberger, 1998; Froemke et al., 2013; Kilgard & Merzenich, 1998; Weinberger, 2007), but the synaptic mechanism underlying the functional plasticity is unknown. To examine how ACh release affects ACx synaptic plasticity *in vivo*, we combined optogenetics with two-photon microscopy. We crossed ChAT-Cre with YFP-H mice (Feng et al., 2000; Rossi et al., 2011) to obtain a double-transgenic mouse line that allows optogenetic stimulation of ACx-projecting BF cholinergic neurons and *in vivo* spine imaging in ACx. In these mice, BF cholinergic neurons express Cre recombinase, and a subset of layer 5 pyramidal neurons (PNs) express EYFP.

BF cholinergic neurons exhibit distinct cortical projection patterns (Kim et al., 2016). To locate the source of cholinergic projections to ACx for optogenetic stimulation, we injected cholera toxin B-488 (CTB-488) into the ACx to retrogradely label ACx-projection neurons (**Figure 1a, b**), and used immunostaining of choline acetyltransferase (ChAT) to label cholinergic neurons. We identified ACx-projecting cholinergic neurons throughout the rostral-caudal axis of BF (**Figure 1c**), concentrating in the ventromedial globus pallidus (GP) and the caudal substantia innominata (SI). The source of projection is confirmed via anterograde tracing, by injecting AAV-Flex-EGFP into the identified ACx-projecting BF region of ChAT-Cre mice (**Figure 1—figure supplement 1**).

**Figure 1.**
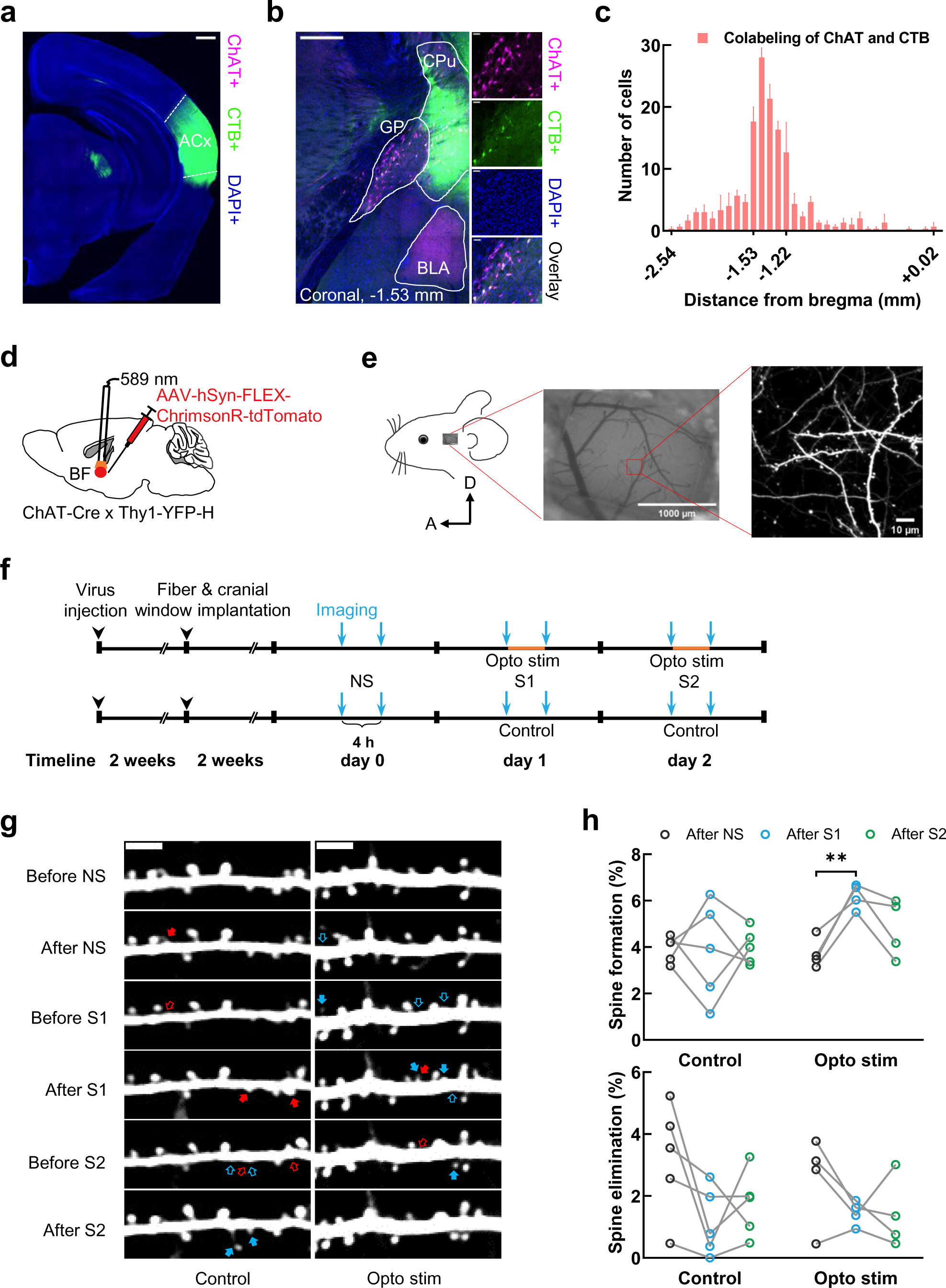
ACh release by optogenetic stimulation of basal forebrain cholinergic neurons induces spine formation in the auditory cortex. **(a)** Image showing CTB injection site in the ACx. Scale bar, 500 μm. **(b)** Immunostaining of ChAT antibodies showing cholinergic (ChAT+) neurons retrogradely labeled by CTB injection in the ACx. CPu, caudate putamen; BLA, basolateral amygdala; GP, globus pallidus. Scale bar, 500 μm (left), 50 μm (right). **(c)** Distribution of labeled ACx-projecting ChAT+ neurons in brain slices along the rostral-caudal axis. n=3, Error bars, s.e.m. **(d)** Schematic showing virus injection and optogenetic stimulation of BF cholinergic neurons. **(e)** Schematic of cranial window surgery and two-photon imaging of spines on the apical dendrites of L5 PNs in the ACx. D, dorsal; A, anterior. **(f)** Experimental timeline for virus injection, fiber and chronic cranial window implantation, and two-photon imaging. NS, no stimulation; S1, stimulation session 1; S2, stimulation session 2. Opto stim, Optogenetic stimulation. **(g)** Example images obtained by repeated imaging of the same apical dendrites of ACx L5 PNs in 2 experimental conditions: Control and optogenetic stimulation (Opto stim). Cyan arrows, newly formed spines. Red arrows, eliminated spines. Scale bar, 4 µm. **(h)** Percentages of spine formation and elimination of apical dendrites of L5 PNs in Control and Opto stim groups after NS, S1, and S2 (Control, n = 5; Opto stim, n = 4. Paired t-test. **P < 0.01).

For optogenetic stimulation of BF cholinergic neurons, we injected AAV-hSyn-FLEX-ChrimsonR-tdTomato virus into the BF of ChAT-Cre × YFP-H mice, and implanted an optic fiber over the injected region for yellow light delivery (**Figure 1d, Figure 1—figure supplement 2**), or crossed ChAT-Cre × YFP-H mice with Ai32 (RCL-ChR2-EYFP) mice (Madisen et al., 2012), and implanted an optic fiber over the BF for blue light delivery (**Figure 1—figure supplement 3**). For *in vivo* observation of synaptic dynamics in ACx, we performed open-skull surgeries over the ACx of ChAT-Cre × YFP-H mice, for repeated high-resolution imaging of spines on the apical dendrites of PNs (**Figure 1e**) (Holtmaat et al., 2009). We quantified the spine formation and elimination rates as a measure of synaptic plasticity as described in previous publications (Knott et al., 2006; Trachtenberg et al., 2002). As spines constantly undergo formation and elimination under baseline conditions, we characterized the baseline formation and elimination rates for each mouse in the absence of cholinergic stimulation by imaging twice at a 4-hour interval on day 0 (**Figure 1f**). On the following day (day 1), we again imaged twice at a 4-hour interval, with or without optogenetic stimulation of cholinergic neurons, as ACh stimulation and control groups, respectively. On the next day (day 2), we performed the same stimulation and imaging experiments as day 1 (**Figure 1f, g**). Through paired comparisons of spine formation and elimination rates for each animal, we found that within 4 hours, the activation of cholinergic neurons increases spine formation in the ACx, without affecting spine elimination (**Figure 1h**), providing direct evidence that ACh release from BF can promote synaptic plasticity in ACx. An additional stimulation session on day 2 did not further increase spine formation (**Figure 1h**), possibly due to limited synaptic protein resource available in PNs.

### Nicotinic AChRs are required for ACh-induced spine formation

Cholinergic receptors consist of two classes, nAChRs and mAChRs, both had been shown to influence synaptic plasticity *in vitro* (Colangelo et al., 2019). To examine which type of receptors function in the observed ACh-induced spine formation *in vivo*, we selectively blocked nAChRs or mAChRs during optogenetic stimulation of cholinergic neurons by either applying a nAChRs antagonist (MEC) or a mAChRs antagonist (Scopolamine) during optogenetic stimulation. For each mouse, we performed optogenetic stimulation of BF cholinergic neurons and two-photon imaging of spines in ACx, as described earlier (**Figure 2a, b**). We found that blocking nAChRs abolished the ACh-induced spine formation, while blocking mAChRs did not have any effects (**Figure 2c**). Note that neither MEC nor Scopolamine affected spine formation rates in the absence of optogenetic stimulation (**Figure 2—figure supplement 1**). These results indicate that nAChRs, but not mAChRs, are required for ACh-induced spine formation in ACx

**Figure 2.**
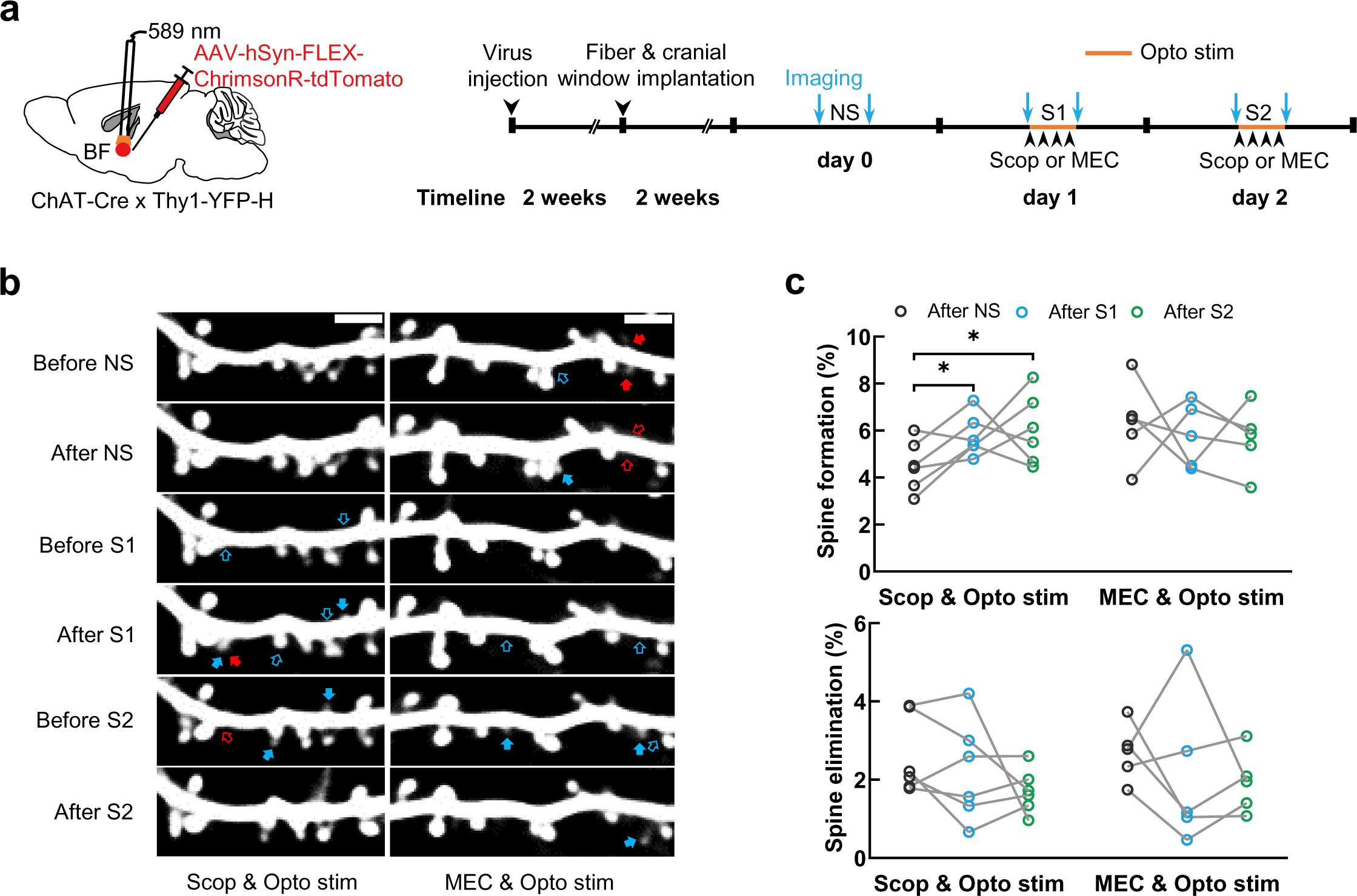
Nicotinic AChRs are required for ACh-induced spine formation. **(a)** Schematic showing optogenetic stimulation and experimental timeline. Scop, Scopolamine. **(b)** Example images obtained by repeated imaging of the same apical dendrites of ACx L5 PNs in 2 experimental conditions: Scop & Opto stim and MEC & Opto stim. Cyan arrows, newly formed spines. Red arrows, eliminated spines. Scale bar, 4 µm. **(c)** Percentages of spine formation and elimination of apical dendrites of L5 PNs in Scop & Opto stim and MEC & Opto stim groups after NS, S1, and S2 (Scop & Opto stim, n = 6; MEC & Opto stim, n = 5. Paired t-test. *P < 0.05).

### GABA transmission is required for ACh-induced spine formation

We observed ACh-induced spine formation on the apical dendrites of cortical PNs, where nAChRs are highly expressed (Colangelo et al., 2019). A previous *in vitro* study had showed that activation of nAChRs increases spine density in hippocampal PNs via a cell-autonomous mechanism (Lozada et al., 2012b). On the other hand, nAChRs are also expressed in cortical GABAergic interneurons, some of which form direct synaptic contacts with the apical dendrites of PNs (Kamigaki, 2019). To determine whether ACh-induced spine formation requires GABAergic connections, we applied a GABA receptor antagonist bicuculline to block GABAergic synapses during optogenetic stimulation (**Figure 3a, b**). We found that bicuculline application abolished ACh-induced spine formation in PNs (**Figure 3c**), indicating that ACh-induced spine formation requires GABA transmission.

**Figure 3.**
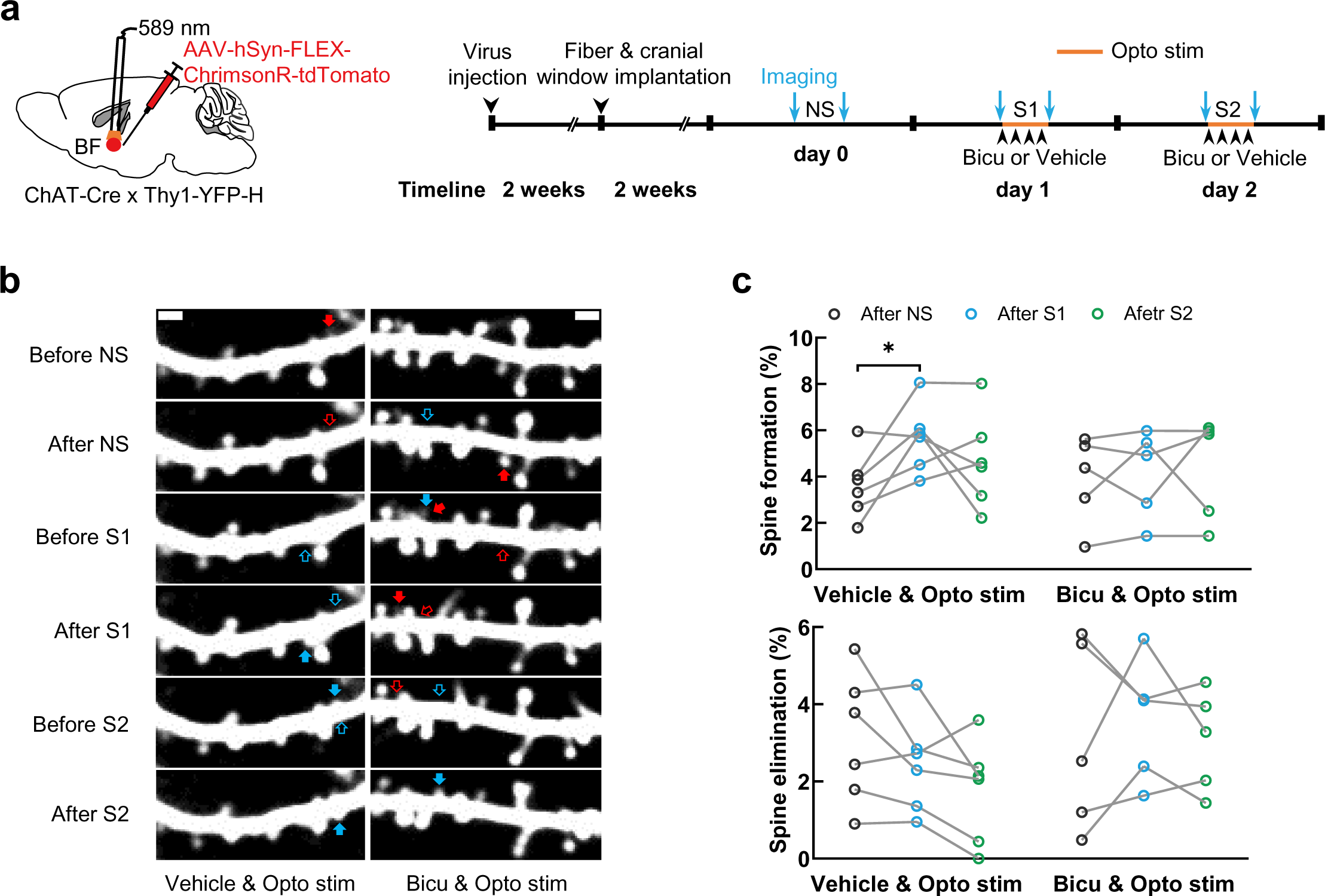
GABA transmission is required for ACh-induced spine formation. **(a)** Schematic showing optogenetic stimulation and experimental timeline. Bicu, Bicuculline. **(b)** Example images obtained by repeated imaging of the same apical dendrites of ACx L5 PNs in 2 experimental conditions: Bicu & Opto stim and Vehicle & Opto stim. Cyan and red arrows, newly formed and eliminated spines. Scale bars, 2 µm. **(c)** Percentages of spine formation and elimination of apical dendrites of L5 PNs in Bicu & Opto stim and Vehicle & Opto stim groups after NS, S1, and S2 (Bicu & Opto stim, n = 5; Vehicle & Opto stim, n = 6. Paired t-test. *P < 0.05).

### Cortical SST interneurons are required for ACh-induced spine formation

To pinpoint which type(s) of GABAergic interneurons in ACx function in the observed ACh-induced spine formation in PNs, we used chemogenetics to manipulate the activity of interneurons in ACx in a cell-type specific manner (Whissell, Tohyama, & Martin, 2016). Among the major types of GABAergic interneurons in the neocortex, somatostatin (SST) and vasoactive intestinal peptide (VIP) expressing interneurons have been shown to be stimulated by ACh (Alitto & Dan, 2012; Fanselow, Richardson, & Connors, 2008; Kawaguchi, 1997; Poorthuis, Bloem, Schak, Wester, de Kock, & Mansvelder, 2013). To specifically inhibit SST or VIP neurons in ACx, we crossed SST-Cre or VIP-Cre mice with YFP-H mice (Taniguchi et al., 2011), and injected AAV-DIO-hM4Di into ACx (**Figure 4a, b**). To target BF cholinergic neurons, we injected AAV-ChAT-hM3Dq, a virus that had been shown to be highly specific for labeling cholinergic neurons (He et al., 2022), into BF. We confirmed the specificity of AAV-ChAT-hM3Dq by staining ChAT antibodies in brain slices injected with the virus (**Figure 4c, d, e**). The ACh stimulation and cortical SST/VIP neuron inhibition was achieved by intraperitoneal injection of CNO, and the efficiency of cholinergic neuron activation and interneuron inhibition was confirmed by c-Fos antibody staining (**Figure 4f, g, h, i**).

**Figure 4.**
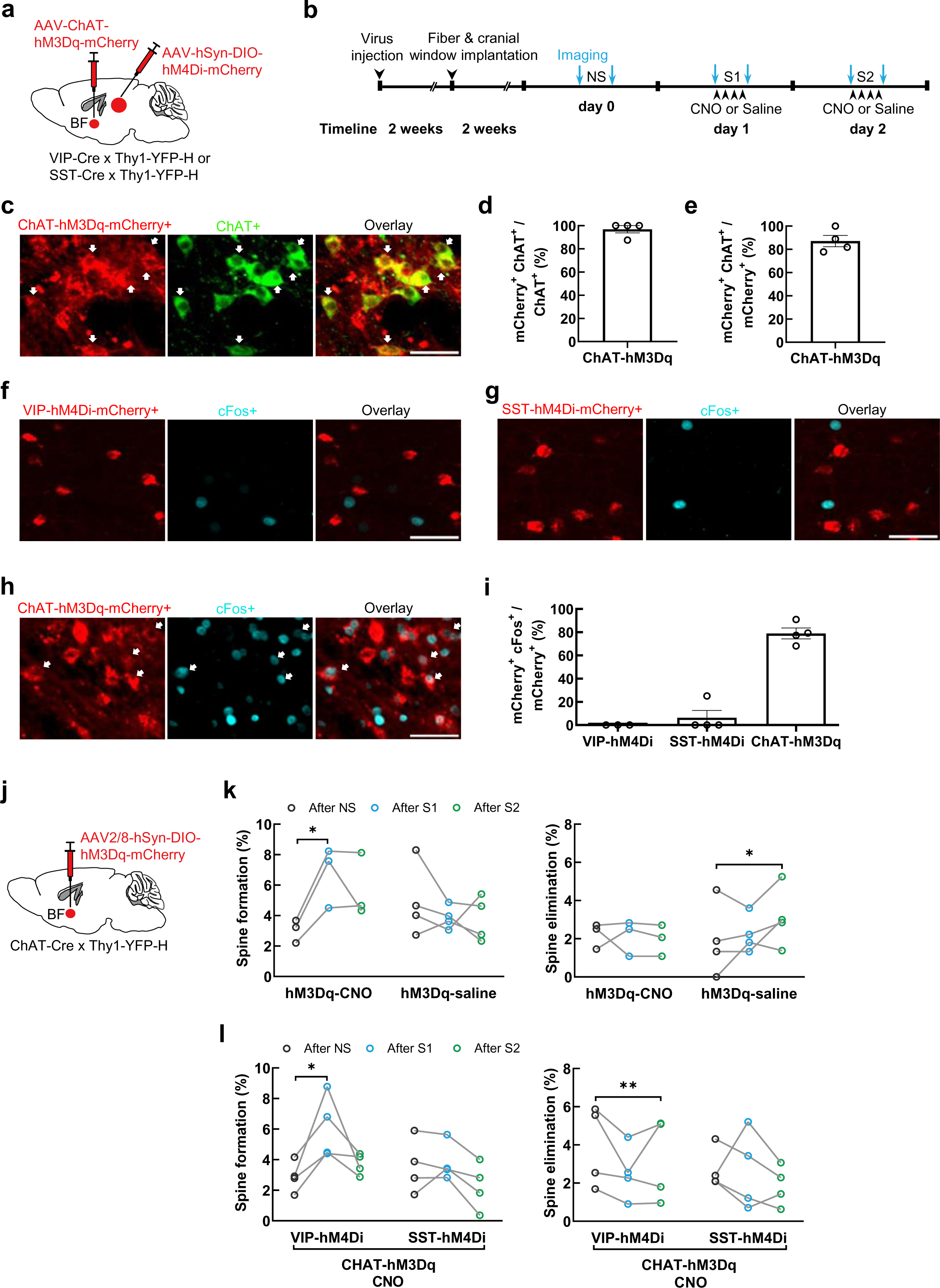
Cortical SST interneurons are required for ACh-induced spine formation. **(a)** Schematic showing virus injection for chemogenetic inhibition of VIP or SST interneurons, and activation of BF cholinergic neurons. **(b)** Experimental timeline. **(c)** Expression of ChAT-hM3Dq in SI & GP cholinergic neurons. White arrows, neurons co-labeled with mCherry and ChAT antibody. Scale bar, 50 μm. **(d)** Percentage of ChAT neurons labeled with mCherry. n = 4. **(e)** Percentage of mCherry^+^ neurons labeled with ChAT antibody. n = 4. **(f, g, h)** Representative images showing co-labeling of cFos with VIP, SST, and ChAT neurons after CNO injection. White arrows point to co-labeled neurons. Scale bar, 50 μm. **(i)** The percentages of VIP, SST and ChAT neurons co-labeled with c-Fos ^+^ after CNO injection. VIP-hM4Di, n = 3; SST-hM4Di, n = 4; ChAT-hM3Dq, n = 4. **(j)** Schematic showing chemogenetic activation of BF cholinergic neurons. **(k)** Percentages of spine formation and elimination of apical dendrites of L5 PNs in hM3Dq-Saline and hM3Dq-CNO groups after NS, S1, and S2 (hM3Dq-Saline, n = 4; hM3Dq-CNO, n = 3. Paired t-test. *P < 0.05). **(l)** Percentages of spine formation and elimination of apical dendrites of L5 PNs in VIP-hM4Di and SST-hM4Di groups after NS, S1, and S2 (VIP-hM4Di, n = 4; SST-hM4Di, n = 4. Paired t-test. *P < 0.05, **P < 0.01).

We validated that chemogenetic stimulation of BF cholinergic neurons could also induce spine formation in ACx, by injecting AAV2/8-hSyn-DIO-hM3Dq-mCherry into the basal forebrain of ChAT-Cre × YFP-H mice, followed by intraperitoneal CNO injection on imaging days 1 and 2 (**Figure 4j, k**). We then performed the two-photon imaging and ACh stimulation experiments while inhibiting SST or VIP neurons in ACx. We found that silencing SST neurons effectively abolished the ACh-induced spine formation, whereas silencing VIP neurons does not have such effects (**Figure 4l**). Therefore, the activity of cortical SST neurons is necessary in ACh-induced spine formation.

### ACh release by chemogenetic stimulation of basal forebrain cholinergic neurons induces spine formation in posterior parietal cortex

ACh not only regulates neuronal activities in sensory cortex, but also affects attentional processing in posterior parietal cortex (PPC) (Bucci, Conley, & Gallagher, 1999; Bucci, Holland, & Gallagher, 1998). To investigate whether ACh could induce synaptic plasticity in cortical regions beyond sensory cortex, we examined spine dynamics in PPC following chemogenetic stimulation of BF cholinergic neurons. We located the source of cholinergic projections to PPC by injecting CTB-488 into PPC for retrograde tracing (**Figure 5a, b**). We found that PPC-projecting cholinergic neurons were dispersed over the whole basal forebrain (**Figure 5c, Figure 5—figure supplement 1**). We injected AAV2/8-hSyn-DIO-hM3Dq-mCherry into BF, and performed two-photon imaging experiments in PPC while stimulating BF cholinergic neurons (**Figure 5d, e**). We found that the chemogenetic activation of PPC-projecting cholinergic neurons could significantly increase spine formation in PPC (**Figure 5f**). Thus, ACh release from BF can also increase spine formation in PNs in PPC, providing a synaptic basis for the Ach-associated enhancement of attentional behavior.

**Figure 5.**
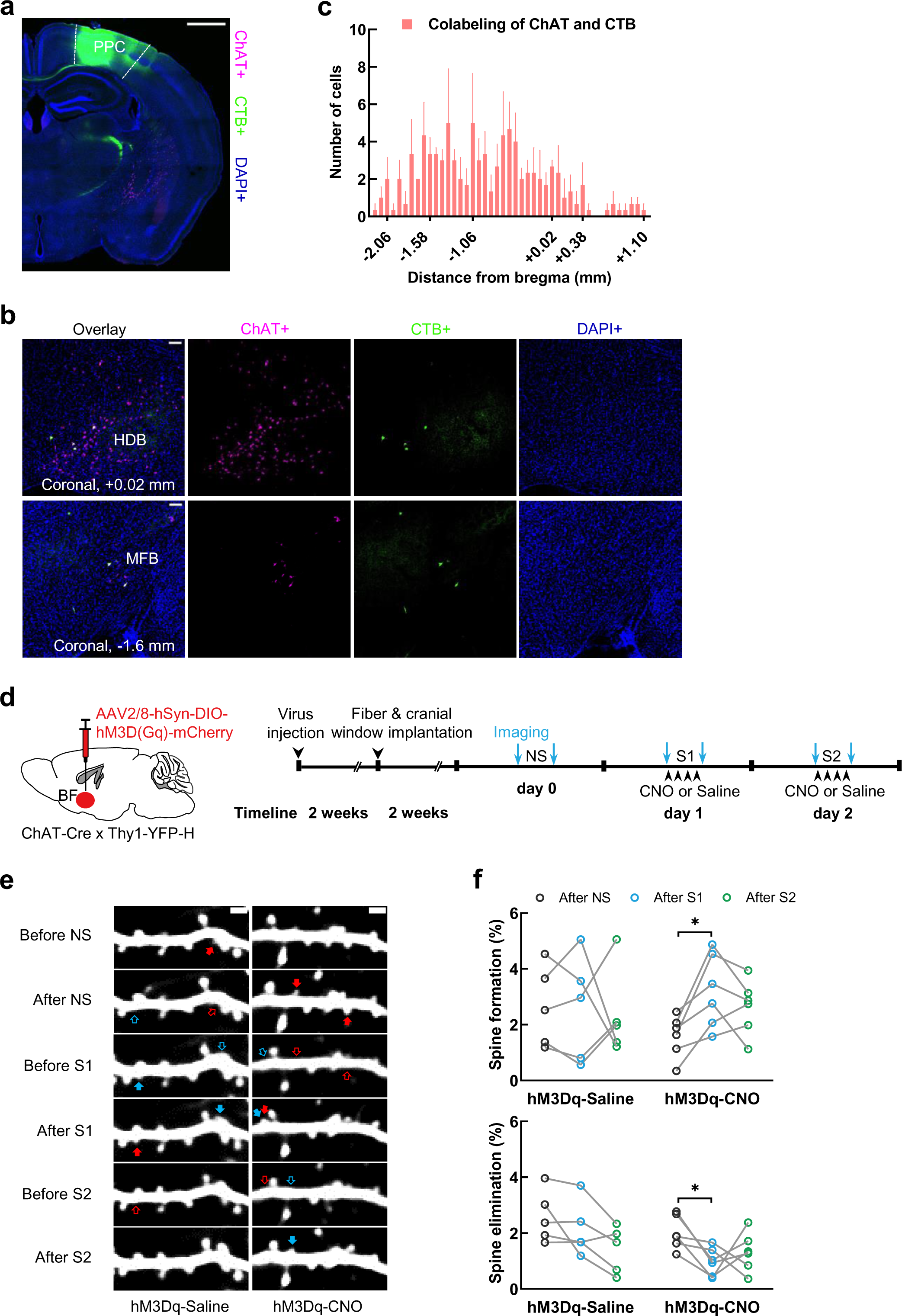
ACh release by chemogenetic stimulation of basal forebrain cholinergic neurons induces spine formation in posterior parietal cortex. **(a)** Image showing CTB injection site in PPC. Scale bar, 500 μm. **(b)** Immunostaining of ChAT antibodies showing cholinergic (ChAT+) neurons retrogradely labeled by CTB injection in PPC. HDB, the nucleus of the horizontal limb of the diagonal band; MFB, medial forebrain bundle. Scale bar, 50 μm. **(c)** Distribution of PPC-projecting ChAT ^+^ neurons along the rostral-caudal axis. n = 3, Error bars, s.e.m. **(d)** Schematic showing virus injection for chemogenetic stimulation and experimental timeline. **(e)** Example images obtained by repeated imaging of the same apical dendrites of ACx L5 PNs in 2 experimental conditions: hM3Dq-Saline and hM3Dq-CNO. Cyan arrows, newly formed spines. Red arrows, eliminated spines. Scale bars, 2 µm. **(f)** Percentages of spine formation and elimination of apical dendrites of L5 PNs in hM3Dq-Saline and hM3Dq-CNO groups after NS, S1, and S2 (hM3Dq-Saline, n = 5; hM3Dq-CNO, n = 6. Paired t-test. *P < 0.05).

## Discussion

Ample evidence from animal studies has showed that BF cholinergic neurons mediate sensory and cognitive functions via ACh release in the neocortex (Bakin & Weinberger, 1996; Bjordahl, Dimyan, & Weinberger, 1998; Bucci, Holland, & Gallagher, 1998; Froemke et al., 2013; Kilgard & Merzenich, 1998; Kim et al., 2016; Lin, Brown, Hussain Shuler, Petersen, & Kepecs, 2015; Meir, Katz, & Lampl, 2018; Parikh et al., 2007; Saper et al., 2010; Teles-Grilo Ruivo et al., 2017; Weinberger, 2007). At the cellular level, ACh has been shown to modulate synaptic plasticity of many types of neurons *in vitro* (Colangelo et al., 2019; Rasmusson, 2000; Xiang, Huguenard, & Prince, 1998). To bridge the knowledge gap between behavioral and *in vitro* studies, we used *in vivo* two-photon microscopy to examine how physiological release of ACh modulates synaptic plasticity in the mouse neocortex. We found that ACh release from BF cholinergic neurons significantly increases spine formation in both ACx and PPC, which requires nAChRs and the activity of SST inhibitory interneurons. Our work provided a synaptic mechanism for ACh-induced functional plasticity in the cortex, and added evidence for the function of local inhibitory connections upon ACh release.

Many studies have linked nAChRs to synaptic structural plasticity in pyramidal neurons (Lozada et al., 2012a, 2012b; Oda & Tanaka, 2014; Sajo, Ellis-Davies, & Morishita, 2016; Schatzle et al., 2011), most of which support the notion that activation of nAChRs promotes synaptic plasticity and dynamics. Consistent with these results, our work showed that inhibiting nAChRs, but not mAChRs, abolished ACh-induced spine formation. Previous work has found that activation of nAChRs increases spine density in pyramidal neurons via a cell-autonomous mechanism (Lozada et al., 2012b). Using GABA receptor inhibitors, we found that although nAChRs are expressed on the apical dendrites of PNs, where we observed the spine changes, GABAergic connections are required. Thus, a multi-synaptic mechanism contributed to the ACh-induced synaptic plasticity.

Previous studies showed that ACh could activate cortical VIP and SST neurons, but not PV neurons (Alitto & Dan, 2012; Fanselow, Richardson, & Connors, 2008; Kawaguchi, 1997; Poorthuis et al., 2013). VIP neurons mainly targeted SST and PV interneurons, and in turn disinhibit the apical dendrites and perisomatic regions of PNs, respectively (Askew, Lopez, Wood, & Metherate, 2019; Chen, Kim, Peters, & Komiyama, 2015; Kamigaki, 2019; Zaborszky et al., 2018). In contrast, SST neurons directly contact distal dendrites of PNs (Chen et al., 2015). Both SST and VIP neurons express nAChRs; by using chemogenetics and SST-Cre and VIP-Cre transgenic mice, we found that silencing SST neurons could abolish ACh-induced spine formation in PNs, but inhibiting VIP neurons did not have any effect. Thus, ACh possibly acts on nAChRs in SST neurons, which, through synaptic contacts on apical dendrites of PNs, regulate the excitation-inhibition balance of PNs to promote spine formation. Our experiments did not exclude the possibility that nAChRs in PNs also participated in this process via cell-autonomous mechanisms. Future studies are needed to elucidate the involvement of the nAChRs on the apical dendrites.

Formation of spines requires the expression and transportation of many synaptic proteins, kinases, and phosphatases including Calmodulin, Calcineurin, Cofilin, Arp2/3, and Actin (Valles & Barrantes, 2022). After an initial ACh stimulation session which triggered significant spine formation, a second stimulation session on the following day failed to increase spine formation. It is possible that BF cholinergic stimulation triggers transportation and insertion of local postsynaptic proteins to form new spines, rather than translation of synaptic proteins, and thus a second stimulation session could not induce spine formation due to limited cellular resources. How acetylcholine recruits these proteins to form new spines remains to be elucidated.

We observed ACh-induced spine formation in both sensory and higher-order cortices, suggesting that the various functional changes governed by ACh release in different cortical areas may share a common synaptic mechanism. Future studies are needed to investigate the functional implications of ACh-induced spine formation, such as its role in sensory processing, learning, memory, and attention.

## Materials and methods

### Animals

All procedures were permitted by the Animal Care and Ethics Committee at the School of Science and Technology, ShanghaiTech University. ChAT-Cre (JAX stock #028861), Ai32 (RCL-ChR2-EYFP) (JAX stock #024109), VIP-Cre (JAX stock #031628), SST-Cre (JAX stock #013044), and YFP-H (Thy1-YFP-H) (JAX stock #003782) transgenic mice were obtained from Jackson Laboratories. The ChAT-Cre mouse line was crossed with the YFP-H line to acquire the ChAT-Cre × Thy1-YFP mouse line. Then the ChAT-Cre × Thy1-YFP mouse line was crossed with the Ai32 mouse line to get the ChAT-Cre × Ai32 × Thy1-YFP-H mouse line. The VIP-Cre or SST-Cre mouse line was crossed with the YFP-H line to acquire the VIP-Cre × Thy1-YFP or SST-Cre × Thy1-YFP mouse line. All mice were housed and bred in a 12/12h light-dark cycle (8 a.m.-8 p.m. light) in the animal facility of the ShanghaiTech with *ad libitum* water and food. Mice used for viral delivery were 5-6 weeks old. Mice used for imaging experiments were 9-14 weeks old. Each mouse was individually housed after fiberoptic implantation.

### Surgery and virus injections

AAV2/9-CAG-FLEX-EGFP (2.3×10^13^ vg/mL), AAV2/8-hSyn-FLEX-ChrimsonR-tdTomato (2.0×10^13^ vg/mL), AAV2/8-hSyn-DIO-hM3Dq-mCherry (2.6×10^13^ vg/mL), AAV-hSyn-DIO-hM4Di-mCherry (3.3×10^13^ vg/mL) were purchased from OBiO Technology Co., Shanghai, China. AAV-ChAT-hM3Dq-mCherry (5.6×10^12^ vg/mL) was purchased from BrainVTA Co., Wuhan, China.

Mice were anesthetized with isoflurane (1%-1.5%) and positioned in a stereotaxic apparatus (Reward Co., Shenzhen, China). Body temperature was maintained at 37°C using a mini-blanket and heating pad during all surgery and imaging process. Viruses were injected using a glass electrode with a tip diameter of 20–25 µm via a small skull opening (<0.4 mm^2^) with a micro-injector (Nanoject3, Drummond Scientific Co., Broomall, United States). Viruses were diluted to 1.0 × 10^13^ or 6.0 × 10^12^ vg/mL, and 50-150 nL/injection site (1nL/s). The stereotaxic coordinates were as follows: ACx (AP: −2.45 mm, ML: −4.4 mm, DV: −1.1 mm); SI & GP (1. AP: −1.25 mm, ML: −2.36 mm, DV: −3.8 mm; 2. AP: −1.45 mm, ML: −2.52 mm, DV: −3.75 mm); PPC (1. AP: −1.95 mm, ML: −1.55 mm, DV: −0.3 mm, −0.55 mm; 2. AP: −2.3mm, ML: −2.55 mm, DV: −0.6 mm; 3. AP: −2.45 mm, ML: −2.75 mm, DV: −0.5 mm); basal forebrain (1. AP: +0.6 mm, ML: −0.52 mm, DV: −4.9 mm; 2. AP: +0.2 mm, ML: −1.0 mm, DV: −5.1 mm; 3. AP: −0.5 mm, ML: −1.7 mm, DV: −4.8mm, −3.75 mm; 4. AP: −1.2 mm, ML: −1.7 mm, DV: −4.8mm, - 4.1 mm; 5. AP: −1.4 mm, ML: −2.4 mm, DV: −3.5 mm).

For optogenetic stimulation of SI & GP, the optic fiber (0.37 N.A., 200 μm diameter, 4.5mm length, Inper Co., Hangzhou, China) was implanted ∼ 200 μm above the coordinates for viral injection, and the stereotaxic coordinates were as follows: AP: - 1.4 mm, ML: −2.45 mm, DV: −3.5 mm.

For chronic cranial imaging window implantation, mice were anesthetized with isoflurane (1%-5%) and positioned in a stereotaxic apparatus. The skin, subcutaneous tissue, and muscle above the ACx were removed to exposed skull with a scalpel. A 1.5 × 1.8 mm^2^ skull over the ACx was replaced by a 1.5 × 1.8 mm^2^ inner layer and a 2 × 3 mm^2^ outer layer cover-glass window. For the cranial window of PPC, a 3 mm diameter skull over the PPC was replaced by a 3 mm inner diameter and 3.5 mm outside diameter circular cover-glass window, the center stereotaxic coordinates were as follows: AP: −2.1 mm, ML: −2.0 mm. The double-layer cover glass was glued via UV-curing glue. A thin layer of super glue was applied to the edges of the cover glass to seal the cranial window. A custom-made head post was attached to the skull by dental cement to fix the mouse head during two-photon imaging. Mice that have completed the cranial window have recovered for at least two weeks before performing imaging experiments.

After the last imaging session, imaging positions were verified by injecting DiI (10−20 nL) into the cortex according to the vascular pattern of the imaging field. Then the mice were perfused immediately using 0.01M phosphate-buffered saline (PBS) and 4% paraformaldehyde (PFA), sliced (∼100 μm) by vibratome (Leica VT1200S, Wetzlar, Germany) after post-fixed overnight to confirm the imaging position and virus expression.

### Retrograde labeling of ACx- and PPC-projecting cholinergic neurons

For retrograde labeling ACx- or PPC-projecting neurons, mice were anesthetized with isoflurane (1%-5%) and positioned in a stereotaxic apparatus. 500 nL CTB−488 was injected into ACx or PPC using a micro-injector (Nanoject3, Drummond Scientific Co.). After allowing 10 days for retrograde transport, mice were perfused using 0.01M PBS and 4% PFA, sliced (∼50 μm) by vibratome (Leica VT1200S, Wetzlar, Germany) after post-fixed overnight. The injection site was confirmed (Figure 1a, Figure 4a). All brain slices containing basal forebrain were dealt with ChAT immunostaining.

### Immunofluorescence

The primary antibodies include: goat-anti-ChAT (AB144P, Millipore, Temecula, CA, United States, 1:1000), guinea pig-anti-cFos (226308, Synaptic system, Germany, 1:800). The secondary antibodies were as follows: AlexaFluor 647-conjugated donkey-anti-goat (A−21447, ThermoFisher Scientific, Waltham, MA, United States, 1:1000), AlexaFluor 647-conjugated donkey-anti-guinea pig (K0040D-AF647, Solarbio Life Sciences, Beijing, China, 1:1000). CNO or Saline treatment mice were perfused ∼4h after the first administration using 0.01M PBS and 4% PFA, sliced (∼40 μm) by vibratome (Leica VT1200S) after post-fixed overnight. Brain slices were permeabilized with 0.4% Triton X−100 for 30 minutes and blocked with 5% bovine serum albumin (BSA)/PBS for 1 h, then incubated with the primary antibody in 0.02% Triton X−100 and 1% BSA/PBS for 48h at 4℃, then washed with PBS 3 × 10 minutes and incubated with DPAI (1:2500) and the secondary antibody in 0.02% Triton X−100 and 1% BSA/PBS for 2h at room temperature, then washed with PBS 3 × 10 minutes. The brain slices were imaged on a confocal microscope (Nikon CSU-W1).

### Optogenetic manipulation

For optogenetic stimulation, the laser was connected to the implanted optic fiber before each imaging session. For ChR2-mediated cholinergic activation, we used a 473 nm laser (LSR473NL-50-FC, lasever etlong Co., NingBo, China) at a power of 5–10 mW/mm^2^ at the fiber tip, for ChrimsonR-mediated cholinergic activation, we used a 589 nm laser (LSR589H-50-FC, lasever etlong Co.) at a power of 5−10 mW/mm^2^ at the fiber tip. The laser transmission frequency is controlled by PulsePal (Open Ephys) with 20Hz, 1s duration, and 20 pulses, each pulse has a 5 ms duration and 45 ms interval, 1200 trials, ∼4h.

### Drug administration

Clozapine-N-oxide (CNO) (3 mg/kg i.p., MedChemExpress, Shanghai, China), saline, Scopolamine hydrobromide (1 mg/kg i.p., MedChemExpress), Mecamylamine hydrochloride (MEC, MedChemExpres) (1 mg/kg i.p.), Bicuculline (5 mg/kg i.p., MedChemExpress) or vehicle (10% DMSO & 90% corn oil) were administered in 4 divided doses at intervals of 1 h respectively during the first and second stimulation sessions (the first administration was 15 min before stimulation).

### Two-photon imaging

Mice were habituated to the environment of two-photon imaging the day before the first imaging session. Mice were anesthetized with isoflurane (1%−2%) and head-fixed using the custom-made head-post. Image stacks were obtained with a step of 0.7 μm from the cortical surface to 100−150 μm deep using a two-photon microscope (Bergamo II, Thorlabs, Newton, NJ, United States) equipped with a water-immersion 25X 1.05 NA objective (Olympus, Kyoto, Japan) and a Mai Tai Deep See Ti: sapphire laser (Spectra-Physics) at 920 nm. The output optical power <40 mW to avoid phototoxicity. A digital zoom of 6 was used.

### Imaging data analysis

We used an open-source image processing toolbox ORCA (Online Real-time activity and offline Cross-session Analysis) (Sheng, Zhao, Huang, & Yang, 2022) to remove the motion artifacts caused by the respiration and heartbeat of mice. After the image registration, all images were analyzed manually on the three-dimensional stacks using ImageJ and blind to experimental conditions. The same dendritic fragments that were clearly recognizable in all 6 imaging sessions were used for analysis. For spine identification, we adopted criteria from (Holtmaat et al., 2009). In short, the length/neck diameter ratio >3 and head/neck diameter ratio <1.2 of dendritic protrusions (length more than 1/3 diameter of dendritic shaft) were defined as filopodia, other protrusions were classified as spines. Spine formation or elimination was based on the comparison of adjacent imaging sessions. Number of spines analyzed for each mouse was > 150.

### Statistics

All statistical analyses were performed using GraphPad Prism. Two-sided paired Student’s t-tests were used for paired comparisons in Figure 1h, Figure 2c, Figure 3c, Figure 4k, Figure 4l, Figure 5f, and Figure 2—figure supplement 1c.

## Acknowledgements

We thank Drs. Yu Xin and Jun-Qian Qi for suggestions on chronic window surgery; the Molecular Imaging Core Facility at ShanghaiTech University for technical support in confocal imaging and two-photon imaging; and the animal facility of ShanghaiTech for their excellent care of mice. This work was supported by grants from the Ministry of Science and Technology of China (2022ZD0204900), Central Guidance on Local Science and Technology Development Fund (YDZX20233100001002), and Natural Science Foundation of China (31970960) to Y.Y.

## Author contributions

X.Z. and Y.Y. conceived the project. X.Z., Y.Z., and L.Y. performed the experiments and analyzed the data. X.Z. and Y.Y. wrote the manuscript.

## Figure supplement Legends

**Figure 1—figure supplement 1.**
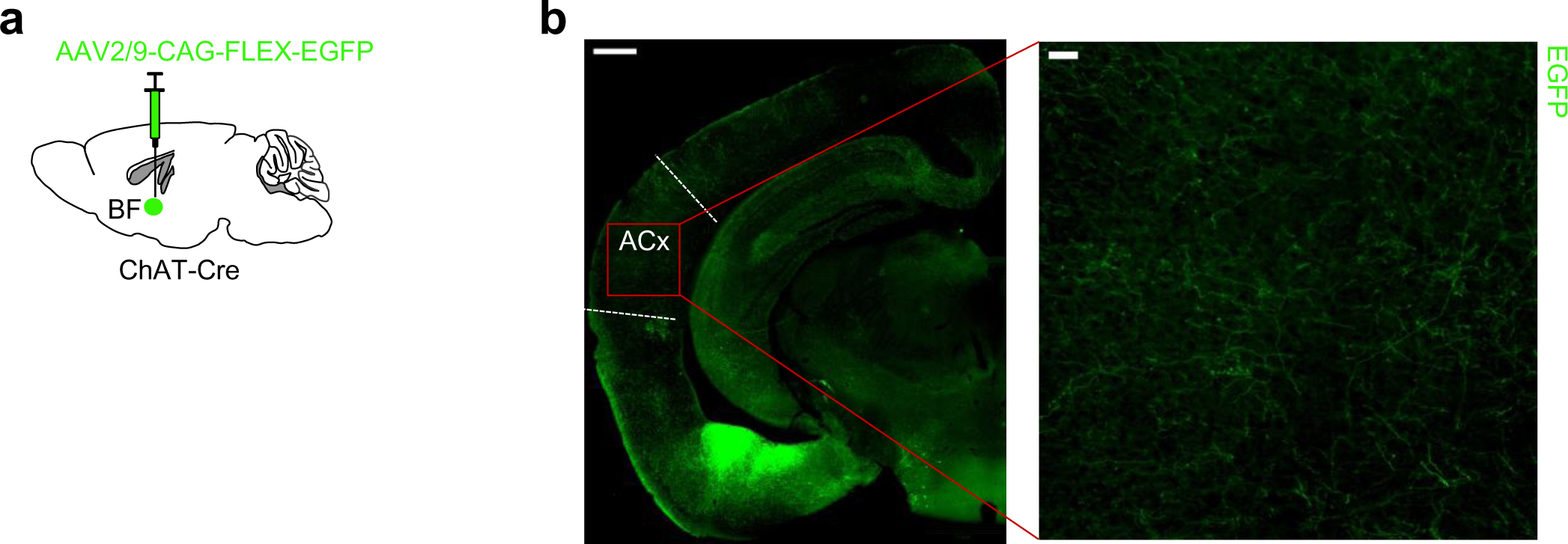
The axonal terminals of cholinergic neurons in the ACx and PPC. **(a)**Scheme of virus tracing of cholinergic neurons in ChAT-Cre mice. **(b)** The axonal terminals of cholinergic neurons are widely distributed in the ACx. Scale bar, 500 µm left, 50 µm right. **(c)** The axonal terminals of cholinergic neurons are widely distributed in the PPC. Scale bar, 500 µm left, 50 µm right.

**Figure 1—figure supplement 2.**
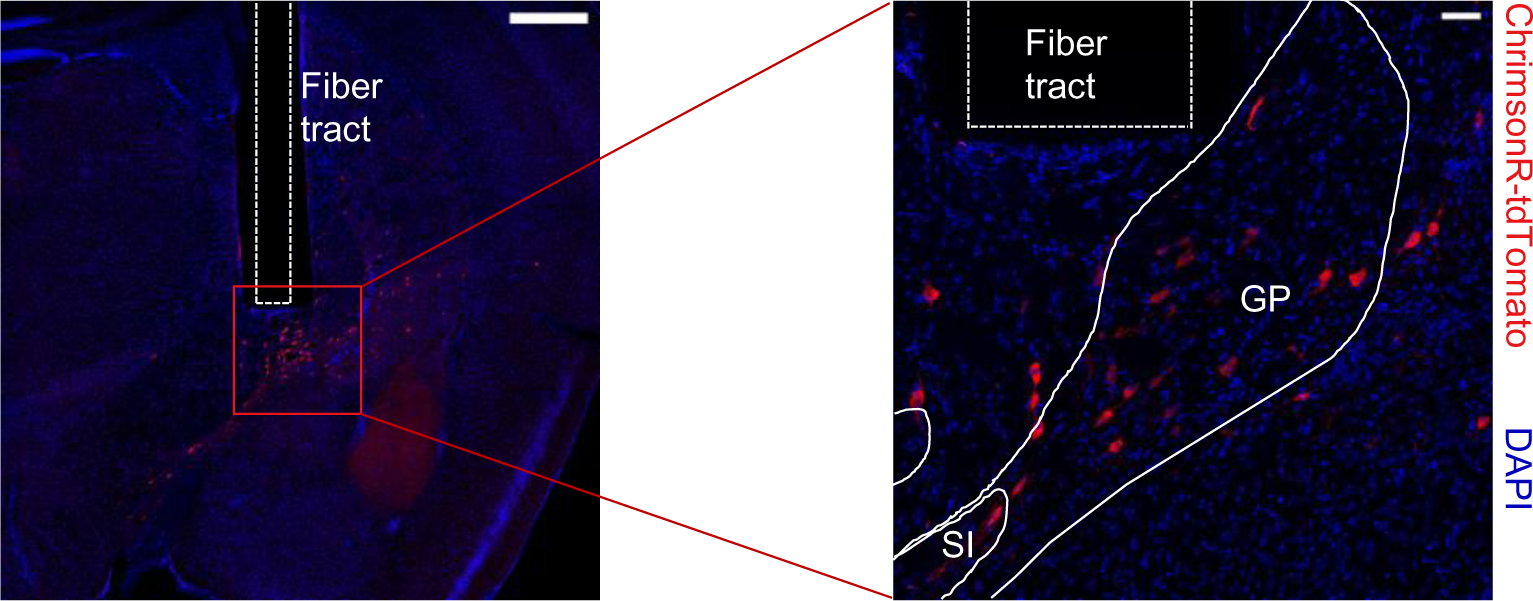
Expression of ChrimsonR-tdTomato in the BF cholinergic neurons. Scale bar, 500 µm left, 50 µm right.

**Figure 1—figure supplement 3.**
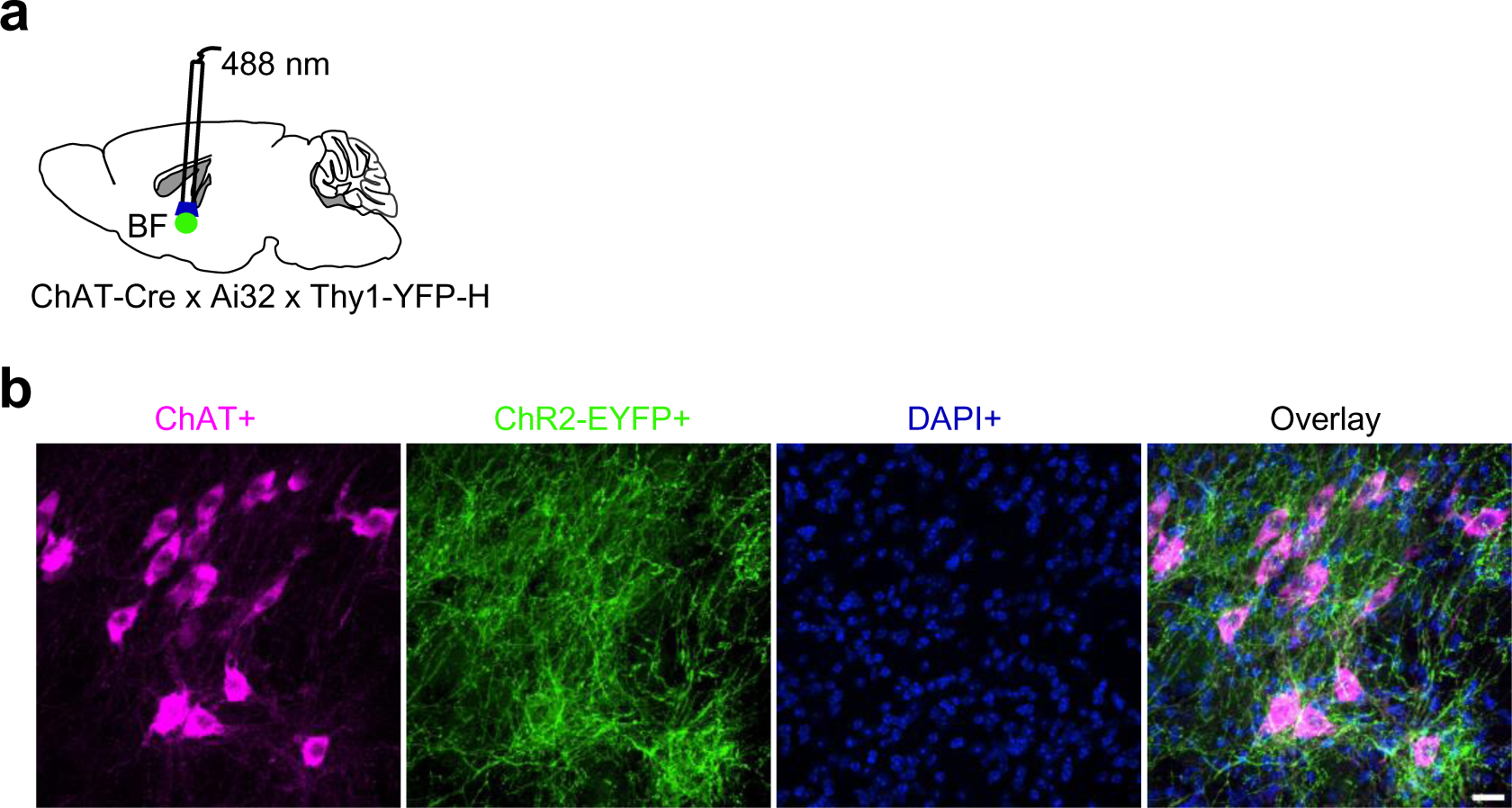
Activation of cholinergic neurons using blue light-sensitive ChR2. **(a)** Schematic showing optogenetic stimulation of BF cholinergic neurons in ChAT-Cre x Ai32 x Thy1-YFP mice. (b) Immunostaining of ChAT antibodies showing cholinergic (ChAT+) neurons co-labeled with ChR2-EYFP in ChAT-Cre x Ai32 (RCL-ChR2-EYFP) mice. For immunostaining, we used ChAT-Cre x Ai32 mice instead of ChAT-Cre x Ai32 x Thy1-YFP mice, because of dense YFP signal in BF of Thy1-YFP mice. Scale bar, 20 μm.

**Figure 2—figure supplement 1.**
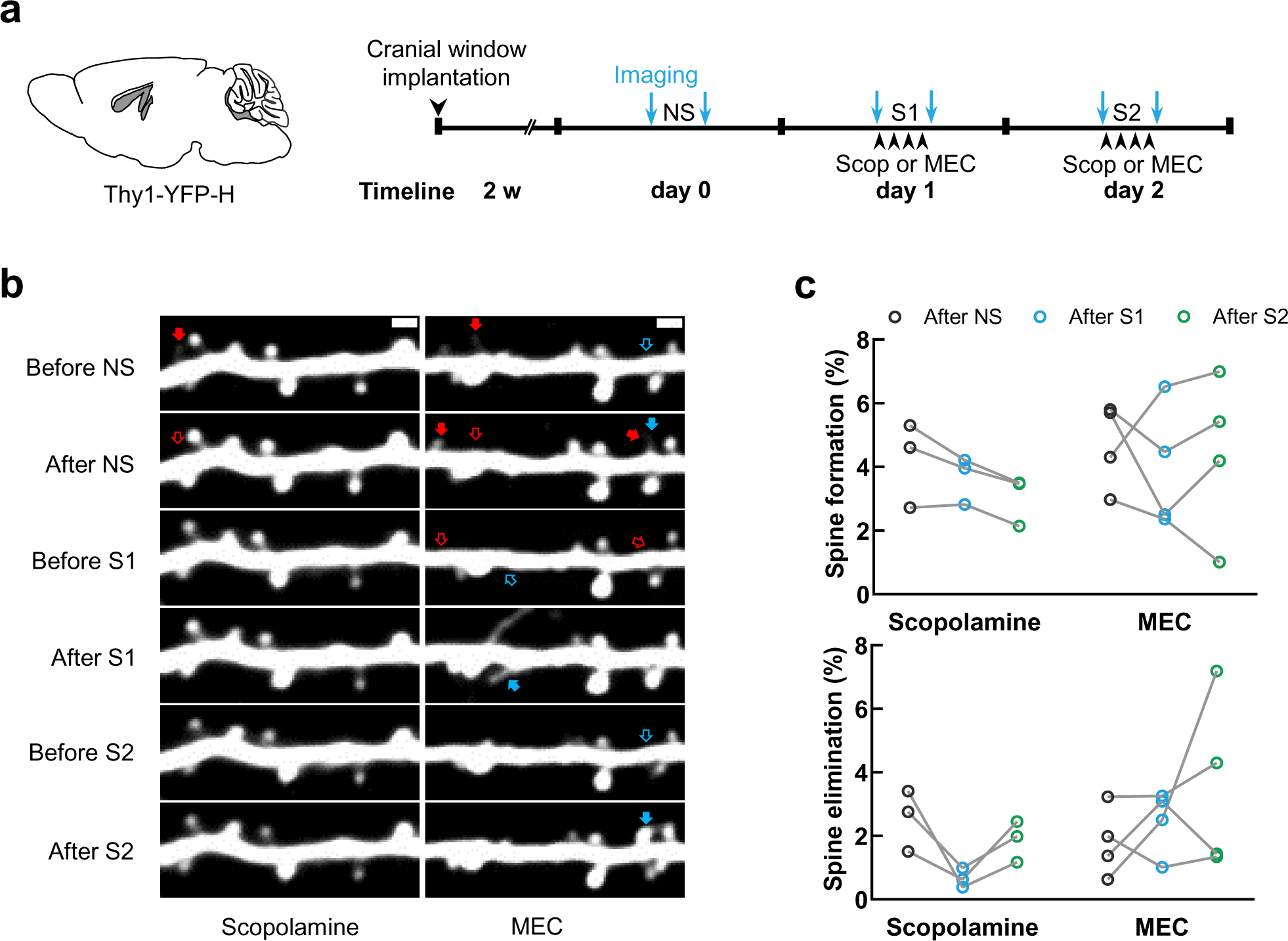
Blocking AchRs does not affect the spontaneous spine dynamics of the ACx. **(a)** Experimental timeline. NS, no manipulation; S1, stimulation session1; S2, stimulation session 2. Scop, Scopolamine. **(b)** Example images obtained by repeated imaging of the same apical dendrites of ACx L5 PNs in 2 experimental conditions: Scopolamine and MEC. Cyan arrows, newly formed spines. Red arrows, eliminated spines. Scale bar, 2 µm. **(c)** Percentages of spine formation and elimination of apical dendrites of L5 PNs in Scopolamine and MEC groups after NS, S1 and S2 (Scopolamine, n = 3; MEC, n = 4. Paired t-test).

**Figure 5—figure supplement 1.**
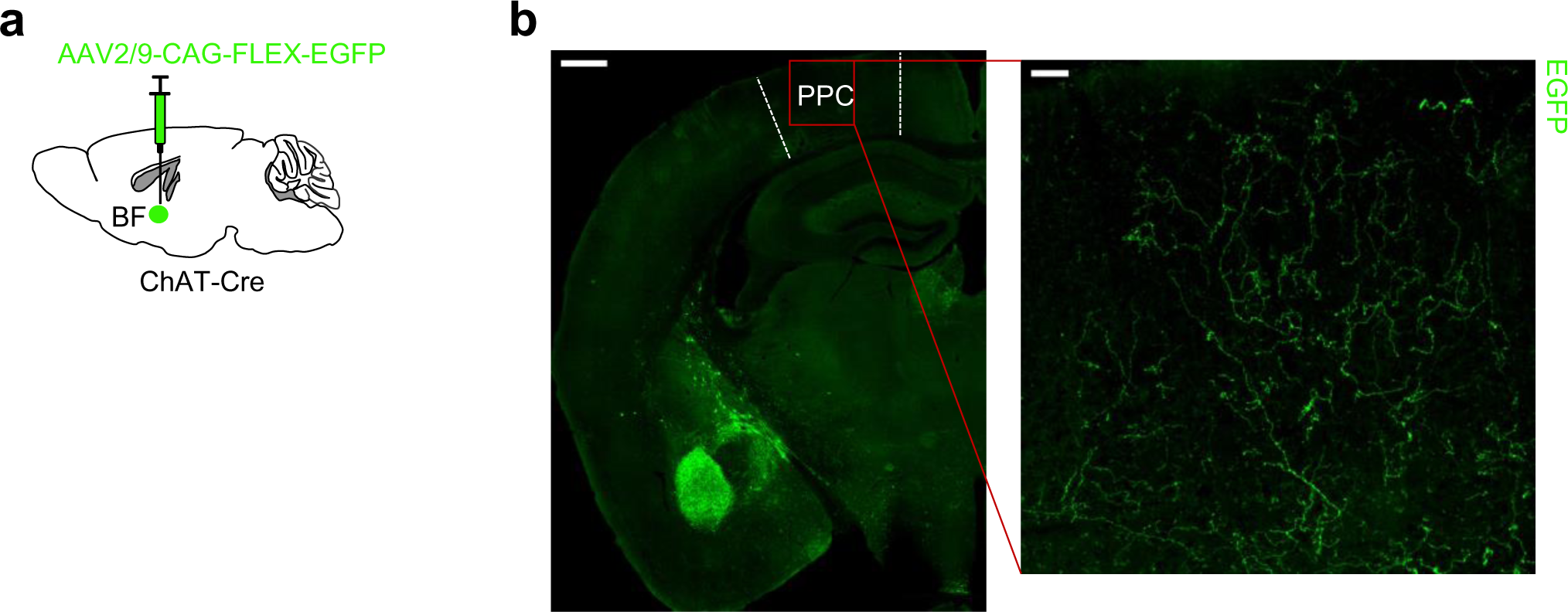
The axonal terminals of cholinergic neurons in the PPC. **(a)** Scheme of virus tracing of cholinergic neurons in ChAT-Cre mice. **(b)** The axonal terminals of cholinergic neurons are widely distributed in the PPC. Scale bar, 500 µm left, 50 µm right.

